# Deficits in olfactory sensitivity in a mouse model of Parkinson’s disease revealed by plethysmography of odor-evoked sniffing

**DOI:** 10.1101/2020.02.27.968545

**Authors:** Michaela E. Johnson, Liza Bergkvist, Gabriela Mercado, Lucas Stetzik, Lindsay Meyerdirk, Patrik Brundin, Daniel W. Wesson

**Affiliations:** Center for Neurodegenerative Science, Van Andel Institute, Grand Rapids, MI, USA; Department of Pharmacology and Therapeutics, University of Florida, Gainesville, FL, USA

**Keywords:** Parkinson’s disease, olfactometer, olfactory deficits, α-synuclein preformed fibrils

## Abstract

Hyposmia is evident in over 90% of Parkinson’s disease (PD) patients. A characteristic of PD is intraneuronal deposits composed in part of α-synuclein fibrils. Based on the analysis of post-mortem PD patients, Braak and colleagues suggested that early in the disease α-synuclein pathology is present in the dorsal motor nucleus of the vagus, as well as the olfactory bulb and the anterior olfactory nucleus, and then later affects other interconnected brain regions. Here, we bilaterally injected α-synuclein preformed fibrils into the olfactory bulb of wild type male and female mice. Six-months after injection, the anterior olfactory nucleus and the piriform cortex displayed a high α-synuclein pathology load. We evaluated olfactory perceptual function by monitoring odor-evoked sniffing behavior in a plethysmograph at one-, three- and six-months after injection of α-synuclein fibrils. At all-time points, females injected with fibrils exhibited reduced odor detection sensitivity, which was detectable with the semi-automated plethysmography apparatus, but not a buried pellet test. In future studies, this sensitive methodology we used to assess olfactory detection deficits could be used to define how α-synuclein pathology affects other aspects of olfactory perception in PD models and to clarify the neuropathological underpinnings of these deficits.

**Highlights:**

- α-synuclein pathology spreads through neuronally-connected areas after bilateral injection of preformed fibrils into the olfactory bulb.
- A plethysmograph and an olfactometer were used for a semi-automated screen of odor-evoked sniffing as an assay for odor detection sensitivity.
- Bilateral olfactory bulb injections of α-synuclein preformed fibrils in female mice led to reduced sensitivity for detecting odors.
- The semi-automated plethysmography apparatus was more sensitive at detecting odor detection deficits than the buried pellet test.

## 1. Introduction

Hyposmia, a reduced sense of smell, affects more than 90% of all Parkinson’s disease (PD) patients, with a minority of these patients experiencing anosmia, a complete loss of their sense of smell (Doty, Deems, and Stellar 1988; Doty 2012). This olfactory dysfunction can occur up to 10 years before the onset of motor symptoms and the clinical diagnosis of PD (Ross et al. 2008; Wu et al. 2011). Currently, the diagnosis of PD heavily relies on the motor manifestations of the disease that correlates with substantial loss of dopamine neurons in the substantia nigra and the widespread deposition of intracellular inclusions, Lewy bodies, and Lewy neurites, partly composed of misfolded α-synuclein fibrils (Postuma et al. 2015). These protein aggregates are able to seed the aggregation of endogenous protein and propagate to interconnected areas (reviewed in (Volpicelli-Daley and Brundin 2018)). According to the Braak staging hypothesis, based on the analysis of post-mortem tissue from PD patients, α-synuclein pathology is initially present in the dorsal motor nucleus of the vagus, as well as the olfactory bulb (OB) and the anterior olfactory nucleus (AON). Later it is present widely throughout the forebrain and eventually it affects the cerebral cortex (Braak et al. 2002; Braak et al. 2004; Del Tredici et al. 2002). The presence of α-synuclein pathology in olfactory structures could explain the olfactory dysfunction seen in PD patients (Beach, Adler, et al. 2009; Beach, White, et al. 2009; Ubeda-Banon et al. 2010).

Animals models of PD-relevant olfactory dysfunction have been established using neurotoxins including, 1-Methyl-4-phenyl-1,2,3, 6-tetrahydropyridine (MPTP) and 6-hydroxydopamine (6-OHDA) (for an extensive review, see (Prediger et al. 2019)). While these models produce neurodegeneration and induce olfactory dysfunction, neither yield robust intraneuronal deposition of misfolded α-synuclein. Genetic models of PD may be used to study olfactory dysfunction, however in these models the olfactory deficits can take a long time to develop (Prediger et al. 2019), and genetic forms of the disease only account for a small percentage of PD cases (less than 10%) (Klein and Westenberger 2012). The mechanism of α-synuclein propagation has been used to reproduce some of the most distinctive hallmarks of PD in mice by the injection of recombinant α-synuclein preformed fibrils (PFFs) (reviewed in (Rey, George, and Brundin 2016). The injection of PFFs into the OB of wild type mice causes Lewy Body-like pathology, which propagates to synaptically connected brain regions over several months (Rey et al. 2016; Rey et al. 2018; Mason et al. 2016; Rey et al. 2019), including key olfactory areas implicated in odor perception. Additionally, these mice have been reported to develop progressive olfactory deficits for odor retention/memory and detection thresholds, using a testing set-up involving a cartridge containing a paper swab impregnated with odor (Rey et al. 2016). In a more recent report, PFFs injections in the OB and/or AON of wild type mice were reported to trigger Lewy Body-like pathology accompanied by sex and age-dependent behavioral deficits, including olfactory dysfunction assessed with the buried pellet test (Mason et al. 2019).

Until now, olfactory deficits have been evaluated in PD animal models using several approaches, such as the buried pellet test, social-odor discrimination test, and odor habituation/dishabituation test (Lehmkuhl, Dirr, and Fleming 2014; Bonito-Oliva, Masini, and Fisone 2014; Zhang, Xiao, and Le 2015; Mason et al. 2019; Rey et al. 2016). While these tests have the advantage of being simple with no requirements for a special apparatus or behavioral shaping, several aspects of them present possible downsides. These can conceptually be synthesized into two main issues: 1) stimulus control, and 2) behavioral monitoring. First, regarding stimulus control, in the assays described above, liquid odor is placed on a substrate (i.e., filter paper, cotton swab), or a food pellet is presented to the animal. While the liquid dispensed onto the substrate is of known intensity, the vapor released is not; thus uncertainty is left in terms of stimulus intensity. Odorant composition of food stimuli is also unstable and may differ considerably from item to item. Second, to monitor behavior in these assays, the observer often manually times and interprets the animal’s response to the odor or food pellet, e.g., when the animal’s snout is within 1 cm of the object containing the odor. While video is often recorded for later scoring, the fact that the animal’s snout is not within a given distance from the odor does not necessarily mean the animal is not investigating the plume (via sniffing). Due to the variability observed using some of the olfactory tests mentioned above, a large number of animals is usually required to detect differences between groups. Taken together, these methods for assaying odor detection and investigation could lead to difficulties in reproducing results and also make it difficult to sensitively define olfactory deficits.

In the present study, we show that bilateral OB injections of PFFs cause olfactory detection deficits in female wild type mice one-, three- and six-months post-PFF injection using a sample size of only 5-6 animals per group. To achieve this, we monitored respiration for changes (or lack thereof) in odor-evoked sniffing which is reflexively displayed by rodents upon detection of a novel stimulus. To monitor sniffing, we adapted a plethysmograph and olfactometer based upon the methods of Wesson et al. (2011) and Youngentob (2005) and delivered animals progressively increasing intensities of odor vapors in an ascending stair-case design. These results, which sensitively and rigorously define odor detection sensitivity deficits following OB injections of PFFs, add to our understanding of the pathological mechanisms of olfactory deficits in PD.

## 2. Methods and materials

### 2.1 Animals

Ten to twelve-week-old male and female C57BL/6N mice were sourced from the Van Andel Institute vivarium. These mice were housed with a maximum of 4 mice per cage under 12-h light/12-h dark cycles with free access to food and water. The housing of animals and all procedures were performed in accordance with the *Guide for the Care and Use of Laboratory Animals* (United States National Institutes of Health) and were approved by the Van Andel Research Institute’s Animal Care and Use Committee.

### 2.2 PD model by bilateral injections of PFFs into the olfactory bulbs

Mouse α-synuclein amyloid aggregates were produced as described in (Volpicelli-Daley, Luk, and Lee 2014) and were kindly provided by Dr. Kelvin Luk, University of Pennsylvania Perelman School of Medicine, USA. Before surgery, PFFs were produced by the sonication of α-synuclein amyloid aggregates in a water-bath cup-horn sonicator for four min (QSonica, Q700 sonicator, 50% power, 120 pulses at 1 s ON, 1 s OFF) and were maintained at room temperature until injection. Mice were anesthetized with isoflurane and injected bilaterally in the OB with either 0.8 µL of PFFs (5 µg/µl; *n*=6 females, *n*=5 males) or 0.8 µL of PBS (phosphate buffered saline) as a control (*n*=5 females, *n*=6 males) (coordinates from bregma: AP: + 5.4 mm; ML: +/- 0.75 mm and DV: - 1.0 mm from dura). Injections were made at a rate of 0.2 µL/min using a glass capillary attached to a 10 µL Hamilton syringe. After injection, the capillary was left in place for three min before being slowly removed. Prior to incision, the animals were injected with local anesthetic into the site of the future wound margin (0.03 mL of 0.5% Ropivacaine; Henry Schein, USA). Following surgery mice received 0.5 mL saline s.c. for hydration, and 0.04 mL Buprenex (Henry Schein, USA) s.c. for analgesia.

### 2.3 Olfactory detection sensitivity testing

#### 2.3.1 Semi-automated system for assaying mouse odor perception

A schematic of the semi-automated olfactory testing set-up, adapted from (Youngentob 2005) can be found in Fig. 1. Product details for the equipment required to create this apparatus are available in Suppl. Table. 1. The concept of the apparatus is to enable monitoring of odor-evoked sniffing mice display upon detection of a novel odor. To accomplish this, two main components need to be in place. First, there need be a method for monitoring respiration/sniffing behavior of the mice. To measure sniffing, we used unrestrained whole-body plethysmography to monitor respiration of the mice as they freely explored the plethysmograph chamber (Data Sciences International). Respiratory transients were detected using a Data Sciences transducer and digitized (0.1-20 Hz) at 600 Hz in Synapse Lite (Tucker Davis Technologies) following a 100X gain amplification (Cygnus Technology Inc). Positive pressure of room air was applied to the chamber using a stable-output air pump (Tetra Whisper). The second main component of the apparatus is an olfactometer allowing precise control of odor delivery. For this, we constructed an odor presentation device (viz., an ‘olfactometer’) adapted from the design of Gadziola et al. (2015). Briefly, custom scripts written in Synapse allowed a user to open one of many relay valves, each when opened allows the flow of clean air into a glass odor headspace vial filled with a few ml of odor (delivered for 6 s duration; odors and liquid dilutions defined below). This then would in effect result in a stream of odor vaporized air (50 mL/min, mixed with 1L/min from the pump) through chemically-resistant Teflon tubing into the plethysmograph chamber. The output from the odor vial was split with a 3-way connector thereby affording the ability to delivery vaporized odor to two plethysmograph chambers simultaneously. Synapse was programmed that upon user initiation of an odor trial, numerous valves would change states which provided acoustic stimuli and possibly other cues to prohibit the mice from associating specific multi-sensory cues with a possible odor and/or becoming aroused by one specific multi-sensory cue and thus influencing their respiration (which is strongly influenced by arousal). Using this set-up, in a longitudinal design, mice underwent olfactory detection sensitivity testing one-, three- and six-months post-PFF injection.

**Fig. 1.**
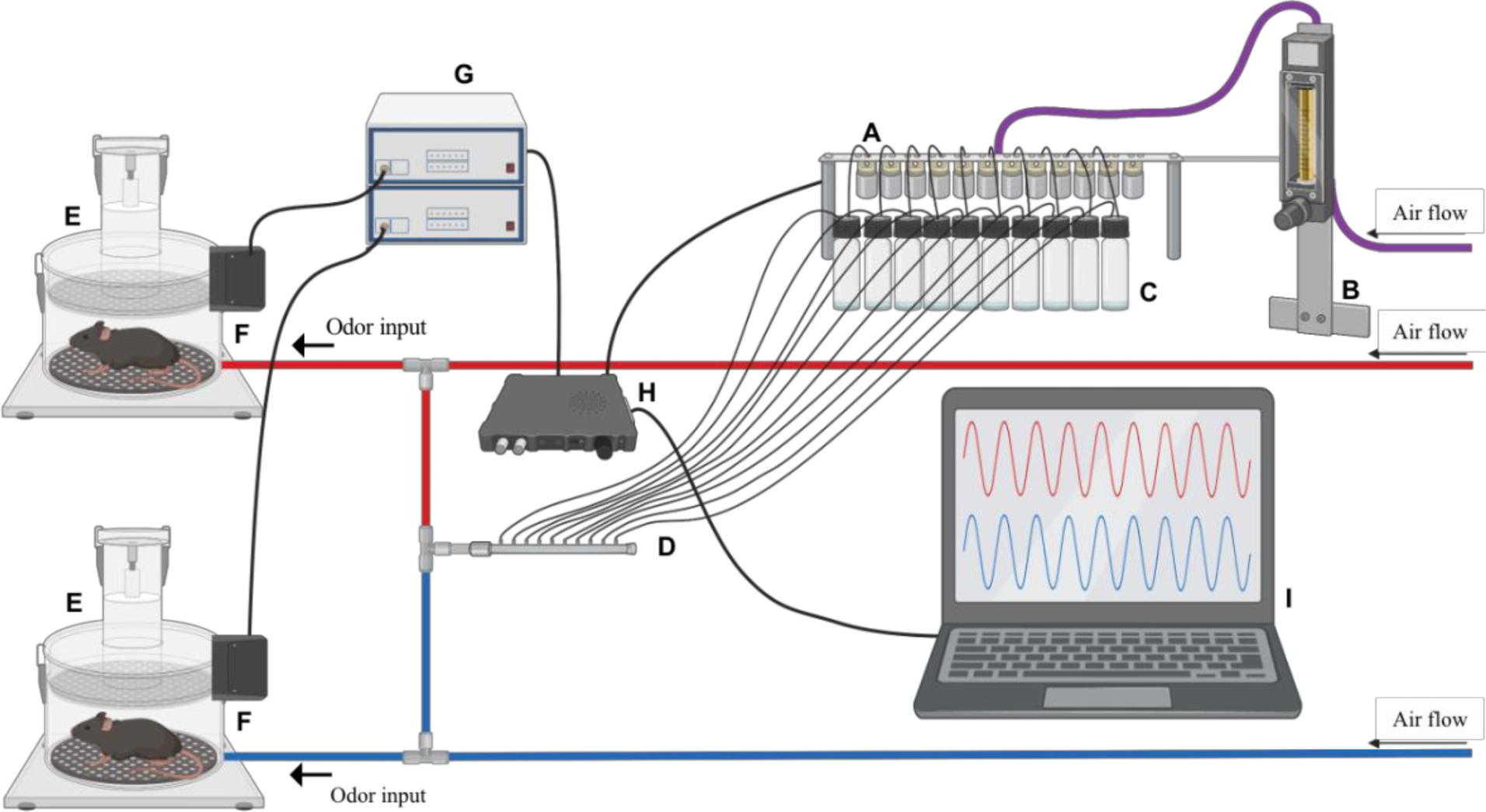
Semi-automated system for assaying mouse odor perception. The olfactometer contains a series of valves (A) that can be digitally triggered through software to open, allowing a set air flow, regulated by a flow meter (B), to pass through the associated valve into the connected odor vial (C). This airflow will continue for 6 s, during which the air flow moves from the headspace vial containing liquid odor through the tubing to combine with the constant air flow (D) to the animals in the plethysmograph chamber (E). The changes in air pressure are recorded through the connected transducer (F). This signal is amplified and filtered (G) before being digitized and stored to the computer (H). Respiration and valve timing are both recorded allowing monitoring of the animal’s respiration in real-time and digital storage (I).

#### 2.3.2 Habituation

Animals were habituated to the testing apparatus for three days prior to testing for each timepoint (months post-injection) so they could acclimate to the plethysmograph chamber, noises and possible mechanosensory/pressure changes which may be associated with valve opening and closing. For habituation the animals were placed in the plethysmograph for 30 min with mineral oil vapor (odor vehicle) presented once per min.

#### 2.3.3 Olfaction investigation testing

Animals were presented with mineral oil once per min for 11 min, then exposed to increasing concentrations of heptanal, starting with 10^−8^ and proceeding in intensity to 10^−2^, also triggered once per min. This protocol was repeated with methyl valerate, isoamyl acetate and 1,7-Octadiene on subsequent days. All odors were diluted with mineral oil to 2 Torr vapor pressure before their serial liquid dilution so that animals’ response to odors could be pooled for analysis.

#### 2.3.4 Analysis

Respiratory data were analyzed in Spike2 software (Cambridge Electronic Design Limited) using a custom script to quantify the percent of time spent in investigatory sniffing after odor-onset. First, common noise rejection was applied to the digital signal from each chamber’s transducer. Following, each signal was digitally filtered (1 Hz to 15 Hz Butterworth, 2^nd^ order), and inspiration peaks were detected by means of identifying the maximum point of each cycle. Instantaneous respiration frequency was calculated off of these peaks and these data were then down-sampled to 100 Hz for ease of handling. The time mice spend sniffing above 6 Hz is considered investigatory behavior (McAfee et al. 2016), as the minimum range of respiration frequency associated with odor investigation in mice is, generally, 6-7 Hz (Wesson et al., 2008). Therefore, off of the instantaneous frequency data, we calculated the duration of time (out of 8 sec of total time following odor delivery) that the animals’ instantaneous respiration frequency exceeded 6 Hz as a measurement of odor investigation. While the odor valve was only open for 6 s, we analyzed an 8 s period as the odor does not immediately diffuse from the chamber. This 8 s period began 1 s after the odor valve opened to allow time for the odor to reach the testing chamber, which was verified in pilot measures using a photoionization detector (Aurora Scientific, miniPID) placed at the input port to the plethysmograph chamber.

#### 2.3.5 Zinc sulfate hyposmia model

As previously described (Lee et al. 2009; Ahn et al. 2018), hyposmia was induced by bilateral intranasal injections with 20 μL of 5% (0.17M) zinc sulfate (Sigma-Aldrich, catalog #7446-19-7) in male C57BL/6N mice. To achieve this, a 25 μL Hamilton syringe (Hamilton, catalog #7654-01) with a 26 G needle (Hamilton, catalog # 7804-03) was fitted with the narrow part of a microloader tip (Eppendorf, catalog # 930001007). The latter provided a long flexible ending allowing easy insertion 4 mm into the naris of the mouse so that the solution could saturate the nasal turbinates. Mice were anesthetized with i.p. ketamine (120 mg/kg; Henry Schein, USA)/xylazine (16 mg/kg; Santa Cruz Animal Health, USA). Once unresponsive, 10 µL of solution was slowly injected over 1 min into the first naris, and the mouse was then immediately inverted (nose pointing down) for 1.5 min to minimize the mouse ingesting the solution, which can cause systemic effects. The mouse was next laid on the injected side for 2 min then on its back for 2 min before the process was repeated with the other naris (Fig. 3A). After both nares had been injected twice (total of 40 µL solution) mice were administered i.p. Antisedan (Santa Cruz Animal Health, USA) 1 mg/kg, to reverse the effects of xylazine. Control mice underwent the same procedure except they received 20 µL of 0.9% saline per naris in place of the zinc sulfate. All mice were allowed 48 h to recover prior to odor detection sensitivity testing in the plethysmograph as described above.

**Fig. 2.**
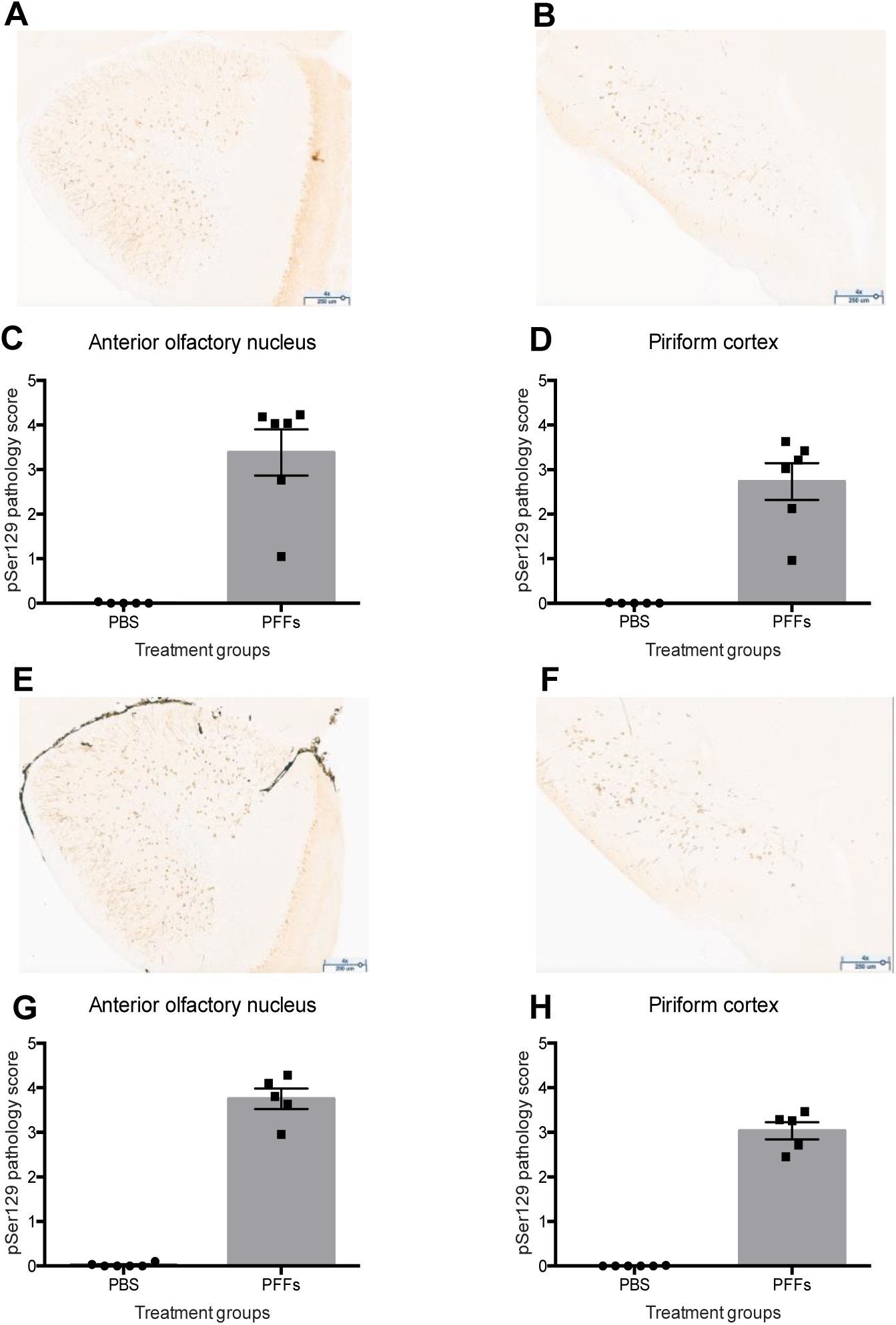
pSer129 α-synuclein pathology in PFFs injected mice. Wild type mice were injected bilaterally in the OB with PFFs or PBS (control). Six-months after injections the load of phosphorylated pathological α-synuclein (pSer129) was assessed by immunohistochemistry. Representative images of the AON and PCx of a PFFs injected female mouse (A & B) and male mouse (E & F) are shown at 4X magnification. The pSer129 immunopositive signal was scored on blinded serial sections including the AON and PCx in female (C & D) and male (G & H) mice. Data displayed as mean ± SEM, n = 5-6 per treatment group.**** = p ≤ 0.0001.

**Fig. 3.**
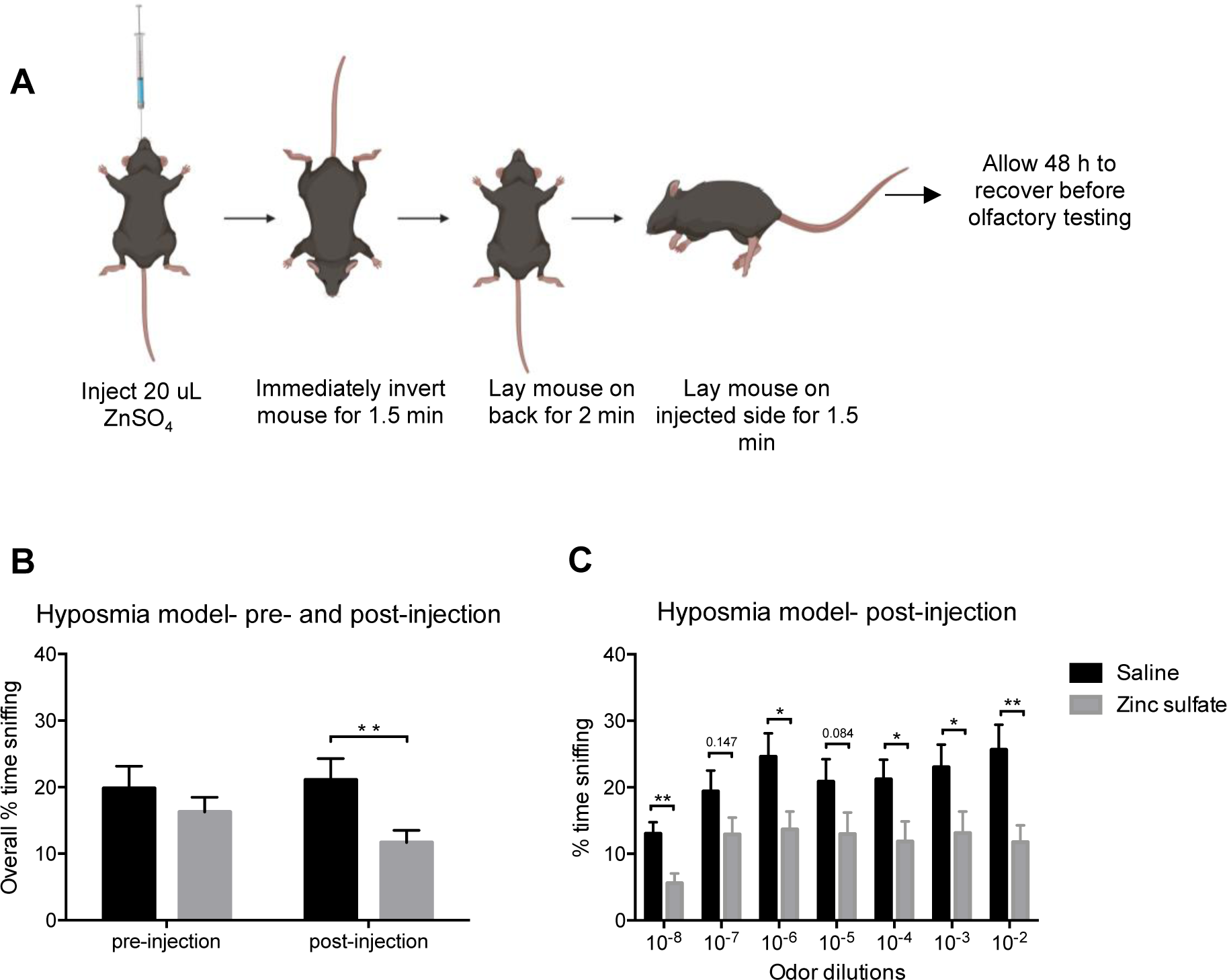
Validation of the semi-automated olfactometer. Schematic describing how the intranasal injections were performed (A). Graphs B & C represent pooled results in response to heptanal, isoamyl acetate, methyl valerate and 1,7-Octadiene odors. Male wild type mice receiving zinc sulfate spent significantly less time engaged in overall investigatory sniffing (B), as well as less time for 10^−8^, 10^−6^, 10^−4,^ 10^−3^ and 10^−2^ odor dilutions compared to the saline group (C). Data displayed as mean ± SEM, n = 9 per treatment group. * = p ≤ 0.05, ** = p ≤ 0. 01.

#### 2.3.6 Statistics

Linear mixed-effects models with square-root transformations were run in R version 3.6.0 to assess overall average instantaneous frequency at each timepoint within each sex. For this analysis random effects for mouse and dilution were used, and odor was included as a covariate. For comparing treatment differences within each odor dilution and sex at each timepoint post-injection (age) we used a linear mixed-effects model that tested instantaneous frequency by treatment adjusted for sex and odor. For both analyses, p-values were corrected for multiple testing using the Benjamini-Hochberg method.

### 2.4 Buried pellet test

The buried pellet test was performed for three consecutive days, with 12 h of fasting in between. Each day of testing, the location of the buried food was changed. The testing paradigm was adapted from Lehmkuhl, Dirr, and Fleming (2014). The buried pellet testing was performed one week prior to the semi-automated olfactory testing described above.

#### 2.4.1 Preparation

Three days prior to the first day of testing, food was restricted for 12 h during the active phase for mice (dark cycle). When the mice had access to food, they were also given one piece of sweetened cereal (Cap’n Crunch) to avoid neophobic food behavior during testing. The starting weight of the animals was recorded prior to food restriction; if an animal lost 10% of its starting weight it was exempt from food restriction and subsequent testing.

#### 2.4.2 Surface pellet test

To control for variability in motivation for food seeking behavior, the surface pellet test was carried out. This was performed as described for the buried pellet test (below), but instead the food item was placed on the surface of the bedding. If a mouse was uninterested in it, it was excluded from subsequent testing. All mice in our study were motivated by this sweetened cereal.

#### 2.4.3 Testing

Animals were moved to individual clean cages containing 2 cm of fresh bedding, covered with a filter top lid, and allowed to acclimate for 5 min. After acclimation, the mouse was put in a clean holding cage while the researcher buried a sweetened cereal item 0.5 cm under the bedding in the testing cage. The mouse was then placed back in the test cage and the latency for the mouse to uncover and put its nose against the food was measured. Once the food was uncovered, the mouse was allowed to eat it. The maximum time allowed to find the food was 5 min; if a mouse was unable to find it within that time period, its latency was recorded as 300 s and it was given the food. Tested mice were not allowed back into their housing cage if it contained untested mice. When all animals in a cage had been tested, they were given free access to food. To avoid outside olfactory cues, the experimenter used double pairs of gloves and changed the outer pair in between mice.

#### 2.4.4 Analysis and statistics

The latency for the mice to uncover the food item was pooled from the three trials and an unpaired Student’s T test was performed in GraphPad Prism version 6.

### 2.5 pSer129 α-synuclein pathology scoring

#### 2.5.1 Tissue preparation

Six-months post-PFF injections animals were anesthetized with sodium pentobarbital (130 mg/kg; Sigma) and perfused through the ascending aorta with isotonic saline followed by ice-cold 4% paraformaldehyde (pH 7.4). Brains were removed, post-fixed overnight at 4°C in 4% paraformaldehyde and subsequently placed in 30% sucrose and store at 4°C until sectioning. Brains were frozen and coronal sections of 40 µm containing the AON and the anterior piriform cortex (PCx) were cut on a sliding microtome (Leica, Germany) and collected as serial tissue sections spaced by 240 µm.

#### 2.5.2 Immunohistochemistry

A series of coronal free-floating tissue sections were stained by immunohistochemistry using a primary antibody directed against pSer129 α-synuclein at 1:10,000 (Abcam, Ab51253) and goat anti-rabbit biotinylated secondary sera at 1:500 (Vector Laboratories, BA-1000). For the detection of the antibody with DAB, we used a standard peroxidase-based method (Vectastain ABC kit and DAB kit; Vector Laboratories). After dehydration, slides were coverslipped with Cytoseal 60 mounting medium (Thermo Fisher Scientific).

#### 2.5.3 α-synuclein pathology *scorings and statistics*

Slides were scanned using an Aperio AT2 scanner (Leica). As previously described (Rey et al. 2016), the serial sections including the entire AON and anterior PCx spaced by 240 µm were scored, in a blinded manner, based on the density of α-synuclein pathology assessed as the load of pSer129 α-synuclein immunopositive signal. Briefly, each section of the region of interest was allocated a value of 0 (no pathology) to 5 (very dense pathology), in increments of 0.5, depending on the pathology load present. The mean score value for each brain region was then calculated for each animal. A permutation test was used to compare the group means of PBS and PFF injected mice for each region of interest.

## 3. Results

### 3.1 pSer129 α-synuclein accumulates in the AON and the PCx

Immunohistological assessment six-months post-PFF injection showed a robust accumulation of phosphorylated, pathological α-synuclein (pSer129) in the AON (females *p* = 3e-07, Fig. 2A & C; males *p* = 1e-07, Fig. 2E & G; 1 synapse from the injection site) and the PCx (females *p* = 3e-07, Fig. 2B & D; males *p* = 1e-07; Fig. 2F & H; 1 or 2 synapses from injection site, direct connection to OB or via the AON) (Cersosimo 2018; Shipley and Ennis 1996; Cleland and Linster 2003). As expected, and consistent with previous literature (Rey et al. 2018; Rey et al. 2016), PBS injected control mice had negligible amounts of pSer129 α-synuclein pathology in these brain regions.

### 3.2 Validation of the semi-automated olfactory test set-up using an established hyposmia model

Rodents display active investigation of novel detected odors upon their perception, which can be quantified by increases of respiratory rate into the stereotyped high frequency respiratory behavior we commonly refer to as ‘sniffing’ (Sundberg et al. 1982; Welker 1964; Wesson et al. 2008; Coronas-Samano, Ivanova, and Verhagen 2016). We reasoned that impaired odor detection sensitivity would be quantifiable by reduced time during odor presentations that mice spend engaged in sniffing when exposed to the increasing odor concentrations. In other words, mice will not display and/or will display less active sniffing for an odor they have yet to detect, but upon detecting an odor for the first time (as in the case for when receiving an odor of sufficiently detectable intensity), the mice will display active sniffing.

To validate that our semi-automated olfactory testing set-up detects reductions in olfactory sensitivity, we used an established method to generate mice that are hyposmic. Thus, we gave wild type mice intranasal zinc sulfate injections, to induce temporary hyposmia (Lee et al. 2009; Ahn et al. 2018), or saline injections as control treatment (Fig. 3A). Prior to intranasal injection, there was no significant difference between mice allocated to the saline or the zinc sulfate group (Suppl. Fig. 1). After intranasal injections, mice exposed to zinc sulfate spent significantly less time engaged in overall investigatory sniffing (9.4% mean difference, *p = 0.003*), indicating hyposmia had been induced (Fig. 3B). More specifically, zinc sulfate-injected mice spent less time engaged in investigatory sniffing for the following odor dilutions, 10^−8^, 10^−6^, 10^−4,^ 10^−3^ and 10^−2^ (Fig. 3C), when compared to the saline group. After verifying the sensitivity of this assay to detect hyposmia, we next use our apparatus to assess olfactory impairments in PBS or PFFs injected mice.

### 3.3 Olfactory detection deficits in mice injected with PFFs

After validating our semi-automated olfactory testing set-up, we explored possible progressive impairments in odor detection sensitivity in wild type female and male mice following bilateral PFFs or PBS injections in the OB (Figs 4A & B). Mice were evaluated in the plethysmograph as above to assess the animal’s odor detection behavior one-, three- and six-months post injection. As a comparison to more traditional methods, all mice were also tested in the buried food pellet test (Lehmkuhl, Dirr, and Fleming 2014). Animals from each treatment group were tested in a pseudo-random manner.

**Fig. 4.**
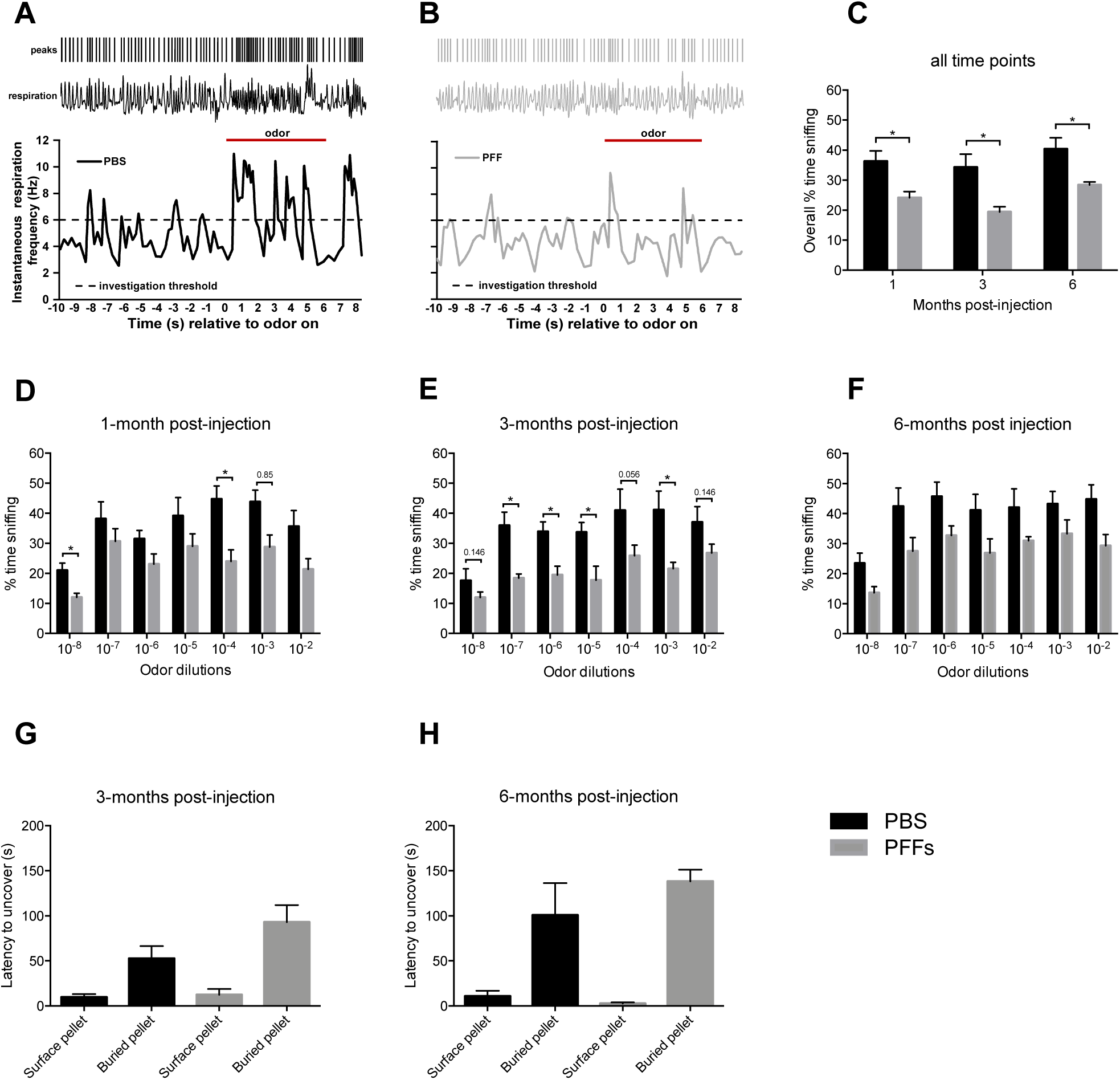
Olfactory deficits in PFF injected mice. Example representative respiration data relative to odor presentation from PBS or PFF treated female mice at three-months post-injection (A & B). Top raster represents inspiration peaks detected offline of digitally acquired respiration signals from the plethysmograph (1 Hz to 15 Hz Butterworth, 2^nd^ order). Instantaneous respiration frequency was calculated off of the inspiration peaks to enable analysis of respiratory frequency dynamics. Horizontal dashed line indicates the threshold operationally defined as ‘investigation behavior’. Odor presentation (6 s) is indicated by the solid horizontal bar (1,7-Octadiene 1×10^−2^ dilution for this example). Graphs C-F represent pooled results in response to heptanal, isoamyl acetate and 1,7-Octadiene odors. Female wild type mice receiving bilateral PFFs OB injections spent significantly less time engaged in overall investigatory sniffing (C) compared to the PBS group at one- (D), three- (E) and six-months (F) post-injection. No difference in olfaction was detected for mice receiving bilateral PFF OB injections using the buried pellet test at three- (G) and six-months (H) post-injection. Data displayed as mean ± SEM, n = 6 for the PFF group and n = 5 for the PBS group. * = p ≤ 0.05.

As expected (Sundberg et al. 1982; Welker 1964; Wesson et al. 2008; Coronas-Samano, Ivanova, and Verhagen 2016), odor delivery elicited vigorous sniffing behavior in mice (Fig. 4A & B). In the examples shown, whereas prior to odor-delivery the mice only occasionally displayed investigatory sniffing (>6 Hz, likely due to spontaneous non-odor driven investigation in the chamber), odor-delivery evoked noticeable increases in sniffing frequency which at least in the PBS-treated animal tended to last for considerable amounts of time during the odor delivery period (Fig. 4A & B). PFF-treated female mice spent significantly less time engaged in investigatory sniffing (Fig. 4C) than the PBS group at one- (12.1% mean difference, *p = 0.024*), three- (14.9% mean difference, *p = 0.018*), and six-months post-injection (11.9% mean difference, *p = 0.018*). This deficit was most prominent at the three-month time point where PFFs injected mice spent significantly less time engaged in investigatory sniffing for the majority of the odor dilutions (10^−7^, 10^−6^, 10^−5,^ and 10^−3^; Fig. 4E). No significant difference was detected between the female PBS and PFFs group at three- and six-months post-injection using the buried pellet test (Fig. 4G & H). At no time point was there a significant difference between PFFs or PBS injected male mice, when evaluated with either the semi-automated olfactory testing set-up (Suppl. Fig. 2A, B, C & D) or the buried pellet test (Suppl. Fig. 2E & F).

## Discussion

We sought to define the influence of bilateral PFFs injections into the OB, and its subsequent pathological accumulation in the olfactory system, on the ability of mice to detect odors. Our results provide new insights into the influence of spreading of α-synuclein pathology on specific abilities of mice to respond to odors at a range of intensities in a behavioral paradigm developed to provide quasi-psychophysical assessments of odor detection sensitivities – a feature reported to be impacted in PD clinical populations (Rahayel, Frasnelli, and Joubert 2012).

Six months after bilateral injections of PFFs into the OB of female and male mice we observed extensive accumulation of phosphorylated α-synuclein in neurons in the AON and PCx. These brain regions are secondary olfactory structures, which receive monosynaptic input from mitral cells and tufted cells located in the mitral cell layer and external plexiform layer of the OB, respectively (Shipley and Ennis 1996; Cleland and Linster 2003). The spread of PFFs from the OB to these and other connected brain structures has previously been well characterized following unilateral injections (Rey et al. 2016; Rey et al. 2018).

In PD, olfactory dysfunction is an early, non-motor symptom. While there are several non-motor signs and symptoms of PD, including mood, gastric, and sleep disorders, olfactory deficits may be more readily tested for, at a low cost, when compared to other non-motor symptoms (Fullard, Morley, and Duda 2017). Olfactory dysfunction has been observed in several rodent models of PD (Kim et al. 2019; Mason et al. 2019; Rey et al. 2018; Rey et al. 2016). However, in the field of neurodegenerative research, olfactory dysfunction in rodents is usually evaluated using systems which are not semi-automated. These methods may have allowed inter-personal differences between observers, such as the interpretation of an animal’s behavior and the observers’ reflexes to manually time specific events, to affect the outcome. We show that our semi-automated plethysmograph-based set-up can detect deficits in odor detection in an α-synuclein-based model of prodromal PD, while the buried pellet test, a manual test, did not detect any differences between groups. Thus, our semi-automated system appears more sensitive, and therefore capable of detecting smaller changes in olfactory perception that may be missed using manual testing. This set up can also be adapted to use ethologically relevant odors, e.g. by replacing the chemical odor with urine from a cagemate vs. an unknown congener.

This is not the first study to use an olfactometer and plethysmograph chamber-based system to assess olfaction in rodents; however, it is the first to apply this type of semi-automated analysis in the context of measuring olfactory impairments in models for PD. Our findings of reduced odor-evoked sniffing upon detection of a novel odor following PFF injections in the OB of female mice substantiates the evidence that α-synuclein pathology in olfactory structures causes olfactory deficits in mice (Rey et al. 2016). Notably, the olfactory impairment was detectable with our apparatus when using only 5-6 female mice per group, which is less than half of the number used previously by Rey et al. (2016). Surprisingly, males did not show any significant olfactory impairment in this study. This is in contrast to Mason et al. (2019) who observed fibril treated males had greater olfactory deficits than females. Sex hormones influence odor perception in rodents (Sorwell, Wesson, and Baum 2008; Wesson et al. 2006) and further investigation into how sex and/or gonadal hormones influence perception in this model is necessary to understand the different olfactory outcomes reported following injections of PFFs.

This new proposed sensitive and rigorous methodology to assess odor sensitivity in PD animal models could be instrumental in determining how α-synuclein pathology affects specific aspects of olfactory perception and help identify the neuropathological underpinnings of these deficits. Future work could assess olfactory deficits following OB injections with a serial dilution of PFFs to correlate olfactory impairment with α-synuclein pathology load, thereby allowing a more in-depth investigation into the relationship supported by the results in this study. Using more sensitive methodology also facilitates the development and evaluation of potential therapeutic interventions aimed to prevent olfactory deficits and the spreading of α-synuclein pathology, as well as the sequential neuronal dysfunction and loss.

## Abbreviations

AON: Anterior olfactory nucleus
OB: Olfactory bulb
PCx: Piriform cortex
PD: Parkinson’s disease
PFFs: α-synuclein preformed fibrils

## Acknowledgements

We thank Emily Schulz for her technical assistance; and the staff of the Vivarium of Van Andel Research Institute for caring for the mice used in this study. We thank Dr. Jennifer Steiner for her helpful feedback on the manuscript. Thanks to Zachary Madaj and Emily Wolfrum from the Van Andel Research Institute’s Bioinformatics & Biostatistics Core for their assistance with pathology and olfactory analysis. The mouse PFFs used in the study were kindly provided by Dr. Kelvin Luk, University of Pennsylvania Perelman School of Medicine, USA. P.B., D.W.W., L.B. and G.M. are supported by the National Institutes of Health (5R01 DC016519-03). P.B. is supported by additional awards (5R37 NS096241-04, M.E.J. and 1R21 NS106078-01A1, L.S.). D.W.W. is further supported by the National Institutes of Health (5R01 DC014443-05).

