## Supplementary files for "Deficits in olfactory sensitivity in a mouse model of Parkinson’s disease revealed by plethysmography of odor-evoked sniffing"

**Supplementary Table 1**

**Suppl. Table 1: Product details for the equipment required to create the olfactory testing apparatus.**

| **item** | **vendor** | **cat#** | **quantity** | **URL** |
| --- | --- | --- | --- | --- |
| #10-32 Male Captivated O-Ring Plug, 5/16" Hex, ENP Brass. Pkg. of 10 | Clippard | 11782-7-ENP-PKG | 1 | https://www.clippard.com/part/11782-7-ENP-PKG |
| 12V 1A power supply | anywhere |  | 3 |  |
| 2-way normally closed solenoid, 12V, 1/8" barb | Parker | 003-0141-900 | 10 | https://ph.parker.com/us/12051/en/series-3-miniature-inert-liquid-valve |
| 3-way normally closed solenoid, 12V, 1/8" barb | Parker | 003-0356-900 | 1 | https://ph.parker.com/us/12051/en/series-3-miniature-inert-liquid-valve |
| 40ml glass headspace vials (case of 100) | Shamrock Glass | 6-01f | 1+ | https://www.shamrockglass.biz/640mlca28mm.html |
| 5 Gal. Black Resin Thermoformed Nursery Pot | Home Depot | Store SKU #520359 | 2 | https://www.homedepot.com/p/5-Gal-Black-Resin-Thermoformed-Nursery-Pot-TFR005G0G18/300434568 |
| 8 Pole Bessel Filter/Amplifier | Cygnus Technology Inc. | FLA-01 | 2 | http://www.cygnustech.com/prices.html |
| Block Manifold #10-32 Threads, 1/16” ID Barbs, 12-Station | Clippard | BTT2-12 | 1 | https://www.clippard.com/part/BTT2-12 |
| Block Manifold #10-32 Threads, 1/8” ID Barbs, 12-Station | Clippard | BTT4-12 | 1 | https://www.clippard.com/part/BTT4-12 |
| BNC cable | anywhere |  | 2 | E.g. Belkin RG58 50-Ohm Thin Ethernet Coaxial Cable with BNC to BNC Male Connectors (6 Feet), https://www.amazon.com/Belkin-50-Ohm-Ethernet-Coaxial-Connectors/dp/B00004Z5KG/ref=sr_1_10?ie=UTF8&keywords=bnc%20cable&qid=1518814622&s=electronics&sr=1-10 |
| Buxco® Small Animal Whole Body Plethysmography | DSI | 601-1425-001 | 2 | https://www.datasci.com/products/buxco-respiratory-products/finepointe-whole-body-plethysmography |
| CED's Spike2 software | CED |  | 1 | http://ced.co.uk/products/spkovin |
| Cole-Parmer PVC Tubing, 1/8" x 1/4", 50 Ft/Pk | Cole-parmer | EW-96605-01 | 1 | https://www.coleparmer.com/i/cole-parmer-pvc-tubing-1-8-x-1-4-50-ft-pk/9660501 |
| Computer/laptop | anywhere |  | 1 |  |
| CUI Desktop AC Adapter, 84W 12V 7A, 2.1x5.5, Level VI | Mouser Elec | 490-SDI90-12-U-P | 1 | https://www.mouser.com/ProductDetail/CUI-Inc/SDI90-12-U-P5/?qs=/ha2pyFadugghadUNUPSuPd176nfrcZNskshP68UsgH51UEjKQoAgQ== |
| Flow transducer w temp and humidity | DSI | 601-2233-001 | 2 | https://www.datasci.com/products/buxco-respiratory-products/finepointe-whole-body-plethysmography |
| Flowmeter, SS float, max. flow rate 0.633 SCFH (299 ml/min) air | Dwyer | va1046 | 1 | http://www.dwyer-inst.com/Product/Flow/Flowmeters/VariableArea/SeriesVA#ordering |
| Hole cap for vials (case of 100) | Shamrock Glass | 6-03d | 1+ | https://www.shamrockglass.biz/60hocaptofit4.html |
| John guest fitting Rigid Elbow, 1/4" Tube OD x 1/8" NPTF Male, 10pk | Amazon |  | 1 | https://www.amazon.com/John-Guest-Acetal-Copolymer-Fitting/dp/B007COMALY/ref=sr_1_1?ie=UTF8&keywords=john%20guest%20fitting%20Rigid%20Elbow,%201/4%22%20Tube%20OD%20x%201/8%22%20NPTF%20Male,%2010pk&qid=1515006824&sr=8-1 |
| LAB RAT EPHYS SYSTEM | TDT |  | 1 | https://www.tdt.com/system/lab-rat-ephys-system/ |
| Male Luer to 500 Series Barb, 1/16" (1.6 mm) ID Tubing | Nordson medical | MLRL004-1 | 1+ | https://www.nordsonmedical.com/Shop/Fluid-Management/Products/MLRL004-1 |
| Male Luer to 500 Series Barb, 1/8" (3.2 mm) ID Tubing | Nordson medical | MLRL013-6005 | 1+ | https://www.nordsonmedical.com/Shop/Fluid-Management/Products/MLRL013-6005 |
| Oversized 440C Stainless Steel Bar 1/8" Thick, 1/2" Wide, | McMaster-Carr | 9575K179 | 1 | https://www.mcmaster.com/9575k179 |
| Polyurethane Ribbon Hose, 1/4” OD-1/8” ID, Multi-Color, 50’ Roll | Clippard | URH8-0804-02T-050 | 1+ | https://www.clippard.com/part/URH8-0804-02T-050 |
| Polyurethane Ribbon Hose, 1/8” OD-1/16” ID, Multi-Color, 50’ Roll | Clippard | URH8-0402-02T-050 | 1+ | https://www.clippard.com/part/URH8-0402-02T-050 |
| Push-Quick Male Compact Connector, 1/4”, #10-32, Pack of 10 | Clippard | PQ-CC08N-PKG | 2 | https://www.clippard.com/part/PQ-CC08N-PKG |
| Stero to bnc adapter | anywhere |  | 1 | E.g. MyCableMart 4 INCH Right Angle 3.5mm Mono Male to Female BNC Adapter Cable, https://www.amazon.com/MyCableMart-RIGHT-FEMALE-Adapter-Cable/dp/B01N51QBRH/ref=sr_1_1?dpID=31e5-SWNQhL&dpSrc=srch&ie=UTF8&keywords=stereo%20to%20bnc%20adaptor&preST=_SX300_QL70_&qid=1518814043&sr=8-1 |
| Teflon septa for headspace vials (case of 100) | Shamrock Glass | 6-04d | 1+ | https://www.shamrockglass.biz/6setetofit15.html |
| Tetra 77851 whisper air pump 10 gallon | Amazon | 77851 | 3 | https://www.amazon.com/Tetra-77851-Whisper-Pump-10-Gallon/dp/B0009YJ4N6/ref=sr_1_1?ie=UTF8&keywords=77851%20tetra&qid=1515007053&s=industrial&sr=8-1 |
| Valve driver board | LabJack | ps12dc | 1 | https://labjack.com/accessories/ps12dc-power-switching-board |
| Marpac Dohm Classic White Noise Sound Machine, White | Amazon |  | 1 | https://www.amazon.com/Marpac-Classic-White-Noise-Machine/dp/B00HD0ELFK/ref=sr_1_6_a_it?ie=UTF8&keywords=white%2Bnoise&qid=1539337504&sr=8-6&th=1 |

**Supplementary Figure 1**

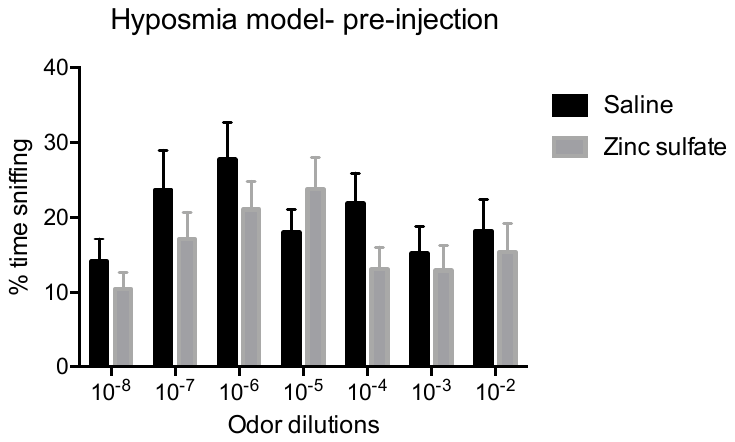

**A**

**Suppl. Fig. 1. Validation of the semi-automated olfactometer.** This graph represents pooled results in response to methyl valerate and 1,7-Octadiene odors. Male wild type allocated to saline or the zinc sulfate group spent a similar amount of time engaged in investigatory sniffing prior to intranasal injections (A). Data displayed as mean ± SEM, n = 9 per treatment group.

**Supplementary Figure 2**

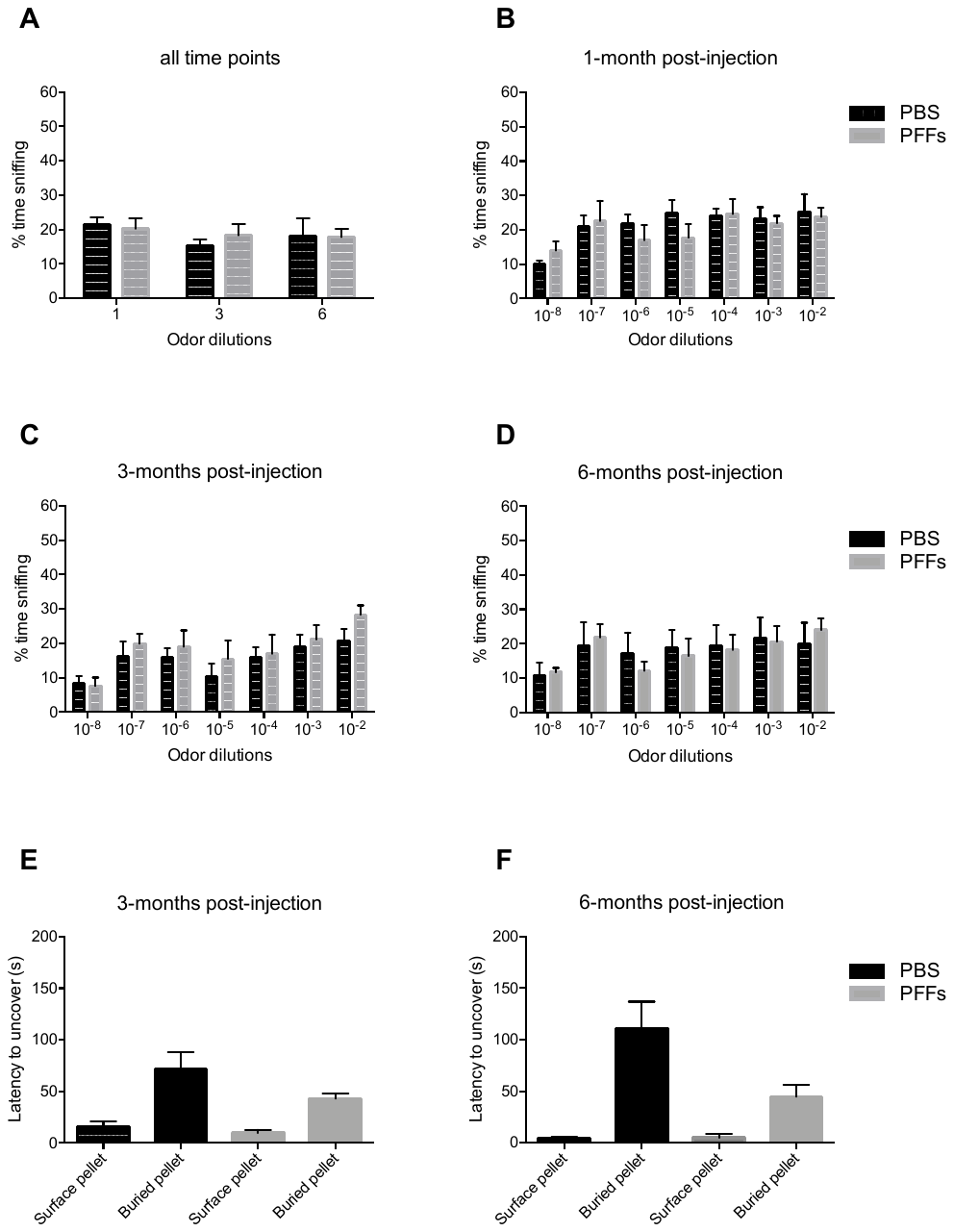

**Suppl. Fig. 2. Olfactory deficits in PFF injected mice**. Graphs A-D represent pooled results in response to heptanal, isoamyl acetate and 1,7-Octadiene odors. Male wild type mice receiving bilateral PFFs OB injections spent a similar amount of time engaged in overall investigatory sniffing (A) compared to the PBS group at one- (B), three- (C) and six-months (D) post-injection. No difference in olfaction was detected for mice receiving bilateral PFF OB injections using the buried pellet test at three- (E) and six-months (F) post-injection. Data displayed as mean ± SEM, n = 5 for the PFF group and n = 6 for the PBS group.
